# Discovery of excited state proton transfer in flavin-based fluorescent protein with large Stokes shift

**DOI:** 10.1101/2025.06.07.658458

**Authors:** Andrey Nikolaev, Elena Markeeva, Eugene G. Maksimov, Anna Yudenko, Elena V. Tropina, Kirill N. Boldyrev, Yuliya Nikolaeva, Dmitry A. Cherepanov, Kirill Kovalev, Yuqi Yang, Valentin Borshchevskiy, Elizaveta Kuznetsova, Alexander Kuzmin, Oleg Semenov, Alina Remeeva, Ivan Gushchin

**Affiliations:** Research Center for Molecular Mechanisms of Aging and Age-Related Diseases, Moscow Institute of Physics and Technology, Dolgoprudny, Russia; Faculty of Biology, Lomonosov Moscow State University, Moscow, Russia; Institute of Spectroscopy RAS, Troitsk, Moscow, Russia; Beijing Institute of Technology (BIT), Zhuhai BIT, Zhuhai, P. R. China; N.N. Semenov Federal Research Center for Chemical Physics, Russian Academy of Sciences, Moscow, Russia; A.N. Belozersky Institute of Physical-Chemical Biology, Lomonosov Moscow State University, Moscow, Russia; European Molecular Biology Laboratory Hamburg, EMBL Hamburg, c/o DESY, Hamburg, Germany; iHuman Institute, ShanghaiTech University, Shanghai, China

## Abstract

Flavin-binding proteins (flavoproteins) are widespread in nature, revealing versatile oxidation-reduction reactions and photochemistry. Flavoproteins derived from LOV domains are used for engineering of ligh-tresponsive tools in optogenetics, as well as fluorescent markers and photogenerators of reactive oxygen species. Despite extensiev efforts, all currently used LOV-derived proteins have similar absorption spectra with maxima around 275, 35-0375, and 450-485 nm. Here, we describe the discovery of a large Stokes shift flavi-nbased fluorescent protein, LSSFbFP, which can be obtained*in vivo* and *in vitro*, with absorption maxima at 340-350 and 395-405 nm. Fluorescence emission of LSSFbFP mirrors that of classical FbFPs with the maximum at ~500 nm. We show that the protein binds lumichrome as the chromophore and use low temperature and time-resolved spectroscopy, X-ray crystallography and modeling to prove that the apparent Stokes shift of LSSFbFP occurs due to excited state proton phenomena observed in flavoproteni s and pave the way for engineering of new flavin-based molecular instruments.

---

Flavins such as flavin mononucleotide (FMN) and flavin adenine dinucleotide (FAD) are ubiquitous cofactors of enzymes and sensory proteins^1^. Flavin-binding proteins (flavoproteins) play many important roles in living organisms, primarily catalyzing oxidation-reduction reactions but also serving as light and redox receptors^2,3^. Versatility of flavins stems from their ability to accept and release electrons and protons, converting between fully oxidized, radical anion, neutral radical, reduced anionic and reduced neutral forms^4^. At the same time, most protein-bound flavins, when not participating in a reaction, remain in fully oxidized form and, due to rigidity of the isoalloxazine moiety, display the same characteristic absorption spectra with maxima around 275, 350-375, and 450-485 nm, imparting them with characteristic yellow color^2^.

While most flavoproteins do not need light to perform their function, some reveal intricate photochemistry^5^. Among the most prominent photoactive flavoproteins are LOV domains, natural blue light receptors^6^. LOV domains were engineered into light-responsive optogenetic tools controlling localization, oligomerization or conformation of the protein of interest^7^. They were also used to engineer flavin-based fluorescent proteins, FbFPs, that have two distinct advantages over GFP-like fluorescent proteins: smaller size of 100-110 amino acids, and independence of fluorescence from oxygen^8^. Finally, efficient LOV-based photosensitizers were also developed^9^.

The major limitation of LOV domain-derived flavin-based molecular tools is the absence of spectral diversity, with all proteins displaying nearly identical absorption and fluorescence emission spectra, which prevents differential activation of LOV-based optogenetics tools or multi-channel recording of fluorescence signals. Recently, we engineered a palette of FbFPs with emission maxima spanning 486-512 nm, which allowed two- or three-color visualization, but required specialized equipment and signal processing.

FbFPs mostly bind FMN, FAD, and riboflavin (RF^11^,) which have almost identical emission spectra. When these flavins are exposed to blue or UV light, they degrade into various compounds, mostly retaining spectral similarity to the original three^12 15^. However, one of the photoproducts, lumichrome (LC), displays markedly different photochemical properties. This difference arises from the complete removal of the ribityl group, which creates an additional titratable site (Figure 1). This modification enables LC to assume different protonation states with distinct spectra. While the definitive list of energetically allowed LC protonation states in aqueous solutions remains a topic of discussion^16,17^, recent comprehensive study provides a compelling insight^18^. At physiological pH levels, LC can exist in both neutral and anionic forms. The neutral form of LC is primarily present in the so-called alloxazinic form and exhibits a blue emission color. The anionic form of LC can be present in N1-anionic and N3-anionic forms, exhibiting green and blue emission colors, respectively. Multiple studies have revealed occurrence of excited state proton transfer (ESPT) in LC in water-alcohol and acidic solvents^19 21^, indicating a decrease in the pK_a_ value of the N1atom following excitation, which could lead to transition from the blue-emitting alloxazinic form to the green-emitting isoalloxazinic or N1-anionic forms. Binding of LC by flavoproteins was cursorily investigated previously, but no distinct phenomena were described^11^.

**Figure 1.**
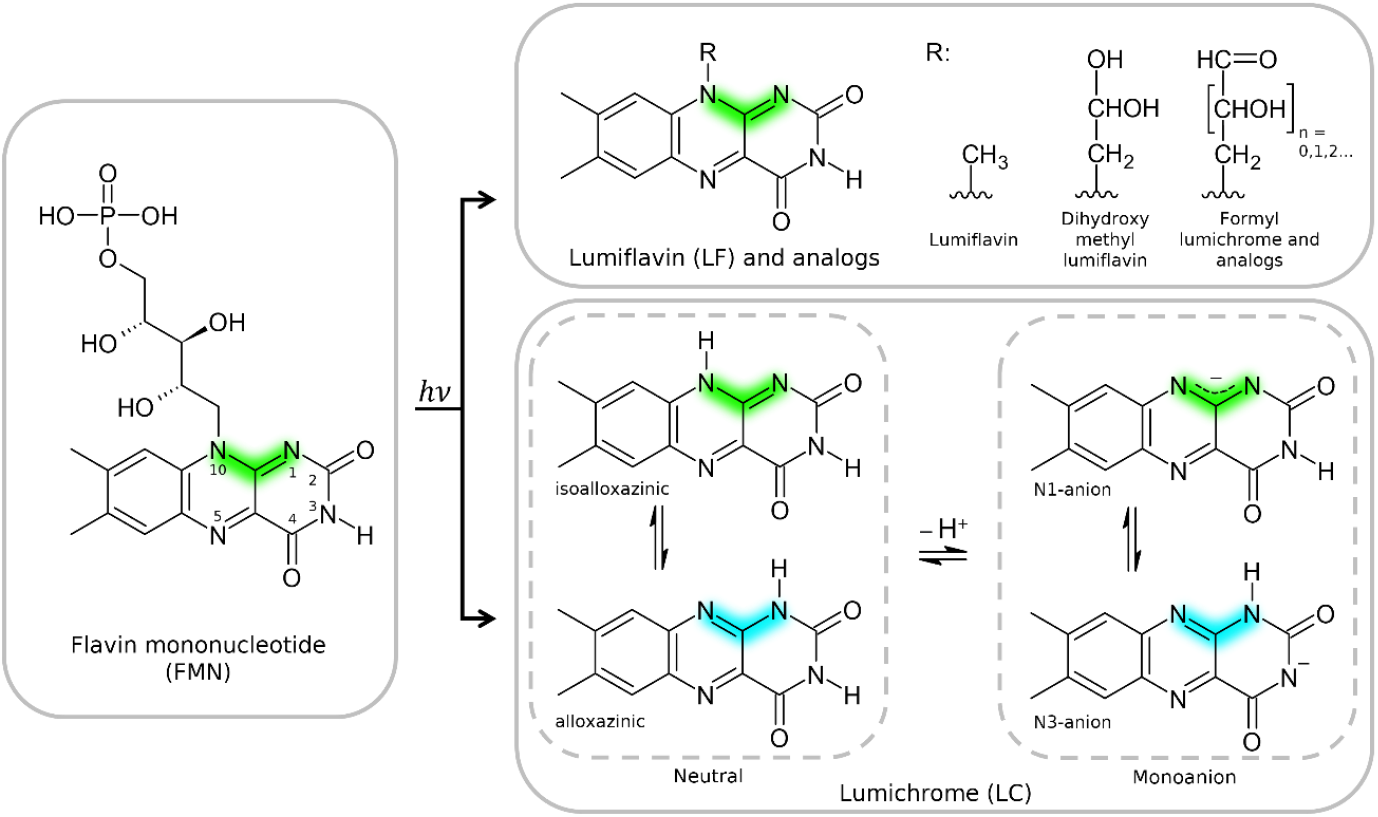
Pathways of FMN photodegradation. In lumichrome, N10 may assume different protonation states, whereas in lumiflavin and analogues it is not titratable. At neutral pH, only the four depicted states of lumichrome are energetically allowed18. Nature of the covalent bonds linking N1 and N10 to the carbon atom C10a defines the energy oeflectronic levels and the expected color of the emitted fluorescence. When N10 and C10a are linked by a double bond, the emission is green, with the maximum around 500 nm, and otherwise it is eb,luwith the maximum around 450 nm13 15.

Here, we not only show that FbFPs can exhibit blue emission in complex with LC but also describe LSSFbFP a novel FbFP that binds LC*in vivo* and *in vitro* and exhibits a substantial Stokes shift of 121 nm, unprecedented for flavoproteins. We investigated the photochemical properties of LSSFbFP using steady state, time resolved and low temperature spectroscopy, obtained structures of LSSFbFP in complexes with FMN and LC, conducted molecular dynamics simulation and carried out DFT calculations to unveil the underlying mechanism of ESPT responsible for its large Stokes shift (LSS).

## Results

### Discovery of large Stokes shift in FbFP

Being interested in development of color-shifted FbFP variants, we began with screening site saturation mutagenesis libraries of CagFbFP, a wel-lcharacterized thermostable flavoprotein^22^. Previously, we tested three of the most promising positions, namely Ile52, Cys85, and Gln148, obtaining a palette of finely tuned variant^1^s^0^. The two latter residues are conserved and involved in signal transduction in wild type LOV domains. In this work, we targeted another conserved position, that of Gln89, which is presumed to not change its conformation during signaling. While most of the examined mutations of Gln89 resulted in modest fluorescence emission shifts, a unique property emerged for the Q89E variant. In contrast to classical flavoproteins, which are distinctly yellow-colored, Q89E variant of CagFbFP appeared completely transpaernt in solution, at the same time exhibiting bright green fluorescence upon excitation with UV light. Its absorption maximum was determined to be located at 390 nm, just on the cusp of the visible spectrum. Due to LSS between the excitation and emission maxima, we dub this variant LSSFbFP thereafter.

Initial attempts at expressing LSSFbFP in*E. coli*, conducted without control of pH and illumination, resulted in inconsistent results. In some cases, the cells were gray-colored, while the protein exhibited LSS, whereas in other cases the cells were yellow-colored and the absorption spectrum of the protein was similar to that of the original protein. This indicated that the LSS was not fully controlled by the protein sequence, which stayed the same, but might have resulted from alterations in chromophore composition due to variations in cultivation conditions. To test this hypothesis, we reconstituted the apo form of LSSFbFP with FMN, RF, FAD and LC chromophores. LSS was observed only for the LSSFbFP-LC complex (Figure S1).

While FAD is synthesized from FMN in *E. coli*, and FMN is synthesized from RF, LC is not normally present in the cells, being a product of flavin photodegradation. To try to control formation of LC *in vivo*, we expressed LSSFbFP in flasks that were either completely covered in black light-proof plastic or wrapped in turned-on LED strips, with pH of the growth medium either controlled (buffered at 6) or not (Figure S2). The protein sample expressed under illumination reproducibly had ~70% molar content of LC, while the sample expressed in the dark had 0% LC. We assume that, similarly to CagFbFP^11^, LSSFbFP binds RF, FMN and FAD*in vivo*, and the exposure to light causes photodegradation of the flavins to LC.

### Biophysical characterization of LSSFbFP

Having obtained samples of LSSFbFP, we proceeded with its characterization, comparing it against the parent variant, CagFbFP. First of all, the mutation Q89E was found to improve the affinity of the protein to LC. The original CagFbFP has a LC dissociaiton constant of 2300 ± 300 nM^11^, while LSSFbFP displays the dissociation constant of 900 ± 300 nM (Figure S3). The dissociation constant of the protein-FMN complex remained essentially the same: 360 ± 50 nM for LSSFbFP versus 350 ± 20 nM for CagFbFP. At the same time, the mutation affected the thermal stability of the complex, shifting the denaturation temperature from 81 °C for CagFbFP-FMN to only 51 °C for LSSFbFP-FMN (Figure S4). In contrast, the thermal stability of the protein-LC complex remained unchanged at 58 °C. The extinction coefficient of LSSFbFP in complex with LC was found to be 8300 M^-1^ cm^-1^ at 350 nm and 6400M^-1^ cm^-1^ at 390 nm. The two proteins have markedly different absorption, fluorescence excitation and emission spectra (Figure 2 and S5). The quantum yield of fluorescence emission of LSSFbFP-LC is 0.28 at 390 nm and pH 8, resulting in the brightness of 1800 M^-1^ cm^-1^. This is lower than the brightness of the original protein in complex with FMN when excited at 440 nm, 5400 M^-1^ cm^-1 11^.

**Figure 2.**
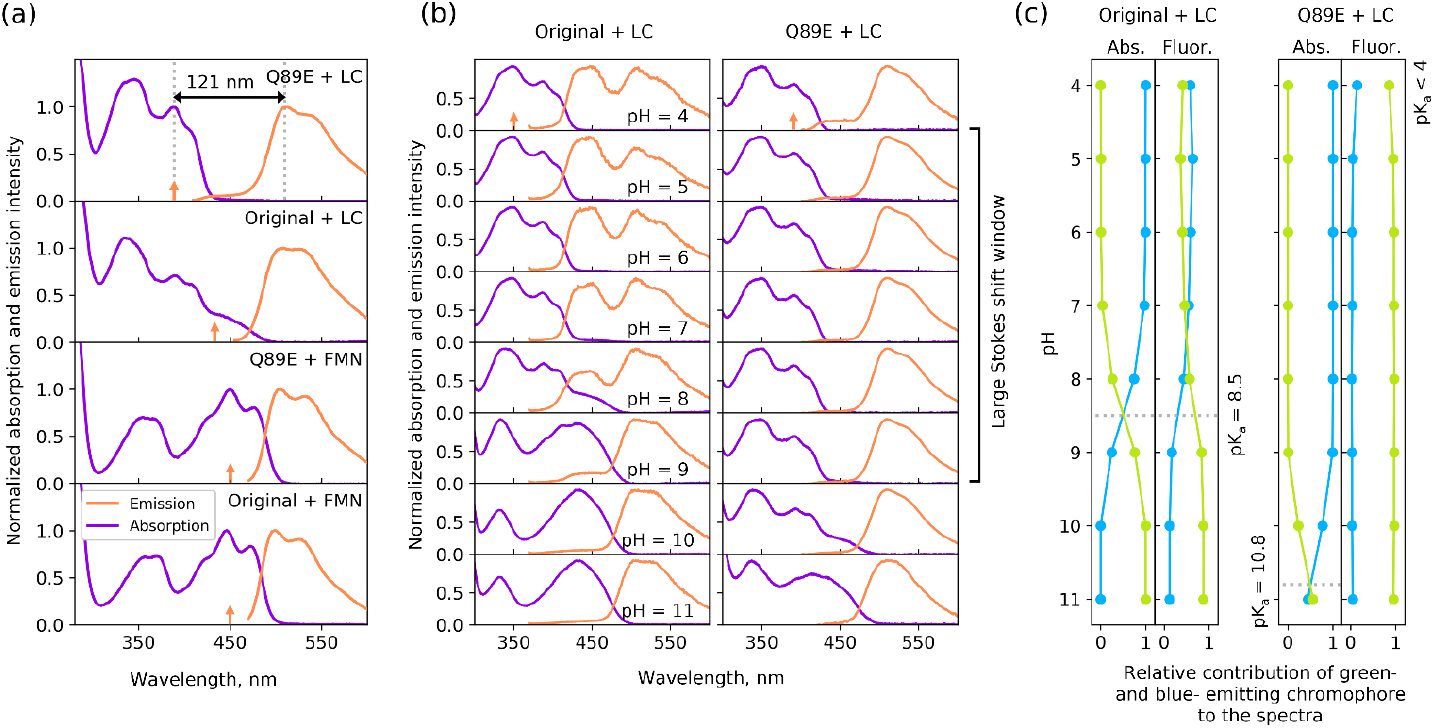
Steady state spectra of CagFbFP and LSSFbFP complexed with different chromophore(s**a.)** Absorption (magenta) and emission (orange) spectra of CagFbFP and LSSFbFP at pH 8. Vertical arrows indicate excitation wavelength. Lumichrom-beound proteins exhibit modified absorption spectra with large Stokes shifts.**(b)** pH dependence of absorption and emission spectra of CagFbFP and LSSFbFP. Excitation spectra at pH 8 may be found in Figure S5**(.c)** pH dependence of relative contribution of different chromophores to theabsorption and emission spectra of protein-LC complexes.

### State of lumichrome in LSSFbFP-LC complex

Given that LC can exist in different protonation states in aqueous solution, it is of great interest to deduce which protonation states are responsible for the absorption and emission of LC embedded in FbFPs. Thus, we measured the pH dependence of the absorption and fluorescence emission spectra of parent CagFbFP and of LSSFbFP in complexes with LC (Figure 2).

At low pH, parent CagFbFP exhibits a blue-shifted absorption, which means that the LC is in its alloxazinic or N3-anionic form. The N3-anionic form seems unlikely, as the amide group of the nearby N117 amino acid should stabilize the hydrogen bound to N3. At high pH, the shape, position, and height of the absorption peaks correspond to the N1-anionic form of LC. Moreover, we can estimate the pKa of the transition from the alloxazinic to the N1-anionic form to be 8.5.

Interestingly, at low pH values, the emission color of the CagFbFP-LC complex is a mix of blue and green components. While the blue component most probably corresponds to the alloxazinic form of LC, the green component can be attributed to an unknown species that appears only after excitation. This indicates the presence of partial ESPT in the parent protein.

Similarly, in the LSSFbFP-LC complex, LC exists in the alloxazinic form at low pH and in the N1-anionic form at high pH. Nevertheless, the pKa value is significantly shifted to the alkaline region and estimated to be 10.8. Contrary to the parent protein, LSSFbFP exhibits emission spectra that correspond to an unknown green-emitting compound, which indicates strong ESPT, essentially at all pH values; only at pH 4 a miniscule admixture of blue-emitting species is observed.

To summarize, in the original protein, LC can exist in the neutral alloxazinic and N1-anionic forms, with blue and green emission, respectively, and a transition pKa of 8.5. While partial ESPT occurs at low pH, it does not lead to LSS. On the other hand, in LSSFbFP, LC exists primarily in the alloxazinic form with blue-shifted absorption, while emitting light from an unknown green-emitting form, causing LSS in a wide pH range (Figure 2.)

### Time dependence of fluorescence emission of FbFP-LC complexes

To gain further insight into temporal evolution of excited FbFP-LC systems, we measured time-resolved emission spectra of both CagFbFP-LC and LSSFbFP-LC at pH 8 (Figure 3a,e). Following the excitation pulse, the emission of CagFbFP-LC exhibited a decrease across all wavelengths, albeit at varying rates (Figure 3b). This implies coexistence of multiple chromophores that are responsible for the fluorescence, with each being characterized by distinct fluorescence spectra and lifetimes. In such instances, employing a global analysis can yield valuable insights. This analytical approach involves fitting the original data using a summation of simple components with mono exponential decays, as represented by the formula:

**Figure 3.**
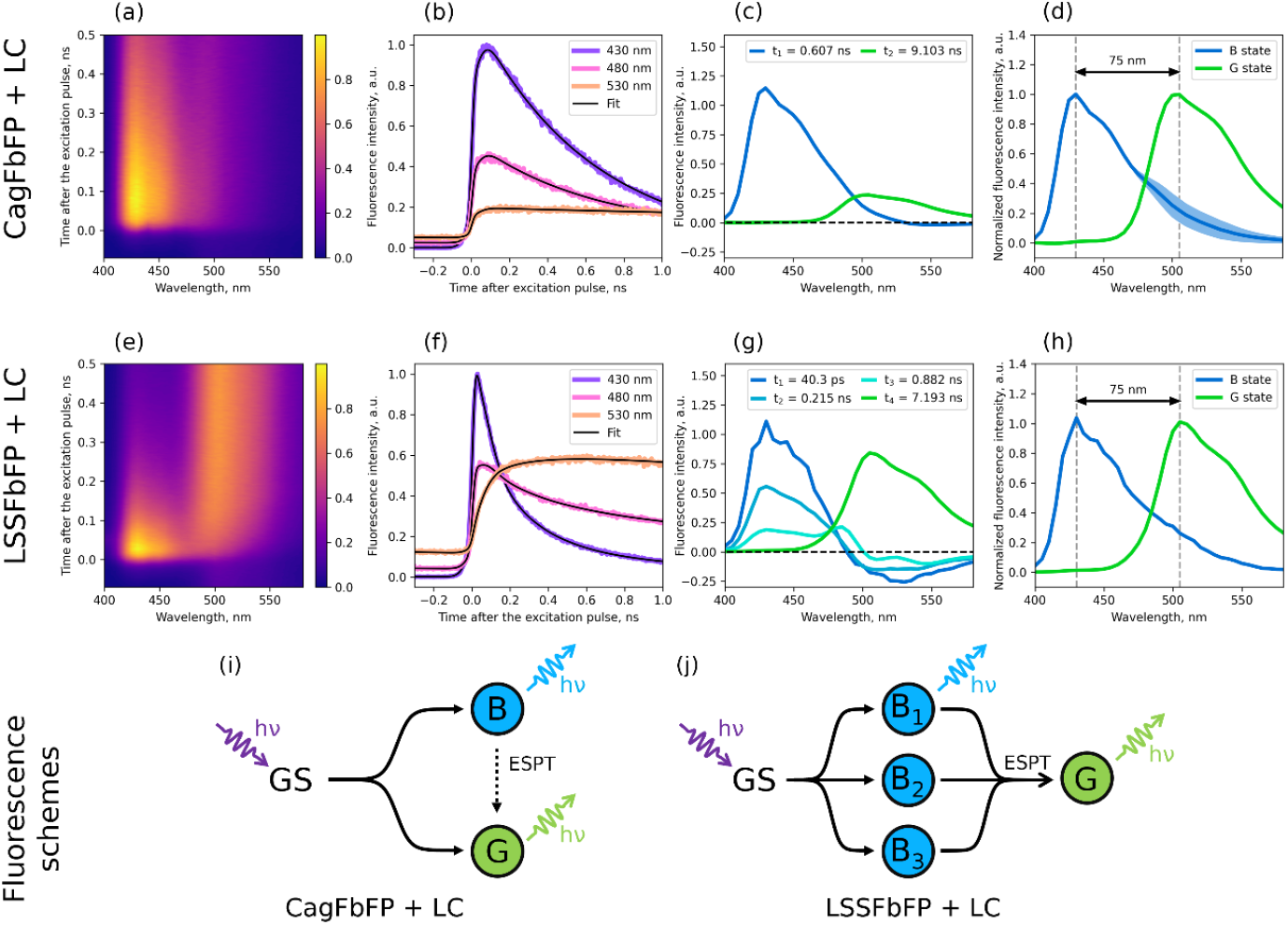
Time-resolved spectroscopy of CagFbFP and LSSFbFP complexed with lumichrome at pH 8**(a**., **e)**Time dependence of emission spectra. **(b, f)** Fluorescence decay traces at different wavelengths.**(c, g)** Decay-associated spectra obtained from global analysis. **(d, h)** Species-associated spectra obtained from global analysis. Species are dubbed B and G due to blue and green emission colo**(i**r,.**j)** Proposed schemes for post-excitation transitions for the LC in complex with CagFbFP and LSSFbFP. Colors of circleisndicate the respective emission colors (B, blue, and G, green).

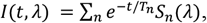

where I(t,) is measured intensity, which is dependent of time t and wavelength, and Sn() and Tn are decay associated spectra (DAS) and lifetimes of respective components. In the case of a solution containing several independent chromophores, global analysis could provide us with the spectra and lifetimes of each chromophore. If components are actively interconverting during the emission process, the DAS can have negative values. In the latter case, DAS can nevertheless be converted to pure species-associated spectra of each component (SAS), albeit with a possible uncertainty.

We conducted global analysis of the time-resolved spectra of CagFbFP-LC, achieving the best fit using two temporal components: a blue, short-lived component and a green, long-lived component (Figure 3c). This is consistent with our previous hypothesis that two forms of LC are responsible for fluorescence. Barely noticeable negative values of the DAS of the blue component indicates weak ESPT to the green form (Figure 3i). SAS of blue and green form can be recovered with a certain degree of uncertainty (Figure 3d).

The time-resolved emission of LSSFbFP-LC unveils a more intricate photophysics. Following the excitation pulse, emission at shorter wavelengths is decaying, while emission at longer wavelengths exhibits an initial increase within the first 0.2 ns, followed by a gradual decay within a nanosecond time frame (Figure 3f). This observation aligns with previous suggestions indicating that LSSFbFP-LC initially absorbs light in its blue-emitting form before swiftly transitioning to its green-emitting form. Global analysis reveals that four temporal components yielded the best fit (Figure 3g). Among these, three decay-associated spectra with the shortest lifetimes shared a similar shape, characterized by a postiive maximum at 425 nm and a negative minimum between 500 and 550 nm. Conversely, the long-lived component exhibited a green-emitting non-negative spectrum with a maximum at 505 nm. Consequently, the blue emission of LSSFbF-PLC demonstrated a fast tri-exponential decay, transitioning to green emission that decayed monoexponentially. The simplest state transition diagram that corresponds to this behavior comprises three blue-emitting states irreversibly transitioning to one green-emitting state post-excitation (Figure 3j).

### Low temperature spectroscopy of FbFP-LC complexes

Further insights into the photophysics of LSSFbFP-LC can be obtained through low-temperature spectroscopy. Lowering the sample temperature significantly hinders or even halts conformational changes within the amino acid environment and slows down chemical reactions as per the Arrhenius equation. We measured the emission spectra of LSSFbFP-LC across temperatures ranging from 300 to 15 K (Figure 4a). The spectrum remained largely unchanged upon cooling to 200 K. However, upon further cooling, the emission peak swiftly shifted towards shorter wavelengths. We assume that the cooling slows down the ESPT and corresponding transition from blue-emitting states to the green-emitting state (Figure 4c,d), thus enabling direct observation of the sharpened emission spectra of the blue-emitting state.

**Figure 4.**
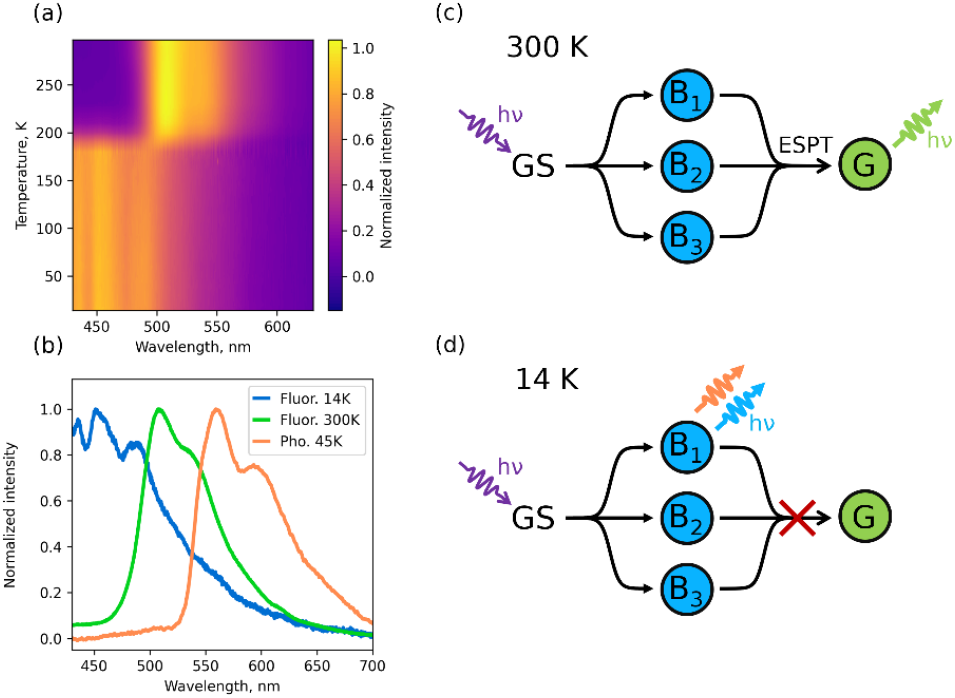
Temperature dependence of LSSFbFP-LC emission. **(a)** Dependence of LSSFbFP-LC fluorescence emission spectra on temperature. **(b)** Fluorescence emission spectra of LSSFbFP-LC at room (300 K) and low (14 K) temperatures and deca-yassociated phosphorescence spectrum at 45 K.**(c-d)** Proposed schemes for post-excitation transitions for the LC in complex with LSSFbFP at different temperatures.Cooling presumably slows down the ESPT.

During the experiments with low temperature spectroscopy, we noticed that when excitation light is immediately turned off, the sample containing LSSFbFP-LC continues to emit yellow light for a fraction of a second. The effect only occurs at temperatures below ~100 K. Similar effect was previously noticed for the original CagFbFP and its variants^23^. We attribute this effect to phosphorescence of the alloxazinic form of LC. Along with ESPT, phosphorescence can be another important factor that lowers the quantum yield of the blue fluorescence of the protein, deepening the gap between emission and excitation spectra. Global analysis applied to time-resolved phosphorescence data allowed us to obtain decay associated spectra of phosphorescence with sufficient signal-to-noise ratio by averaging results from multiple independent experiments (Figure 4b). The estimated phosphorescence lifetime was 125 ms.

### Crystallographic structures of LSSFbFP-LC and LSSFbFP-FMN

We proceeded with our study of LSSFbFP by determining its crystallographic structures in complexes with FMN and LC (Figure 5; also see electron density maps in Figure S6). The structure of LSSFbFP-FMN was determined at pH 6.5 and was essentially identical to the structure of the original CagFbFP^22^ in all regards except for the side chain of 89th position. Root mean square deviation of backbone atoms was 0.2 Å. One of the solvent molecules (MES) crystalized alongside the protein in close proximity to the E89 side chain of one of the protein chains in the unit cell (chain A). Another protein chain, B, displays an alternative conformation of the C terminus, where it is partially dissociated from the remainingβ-sheet, somewhat similar to what is observed in Q148 mutants of CagFbF^2^P^4,25^. Based on analysis of the chain B, the carboxylate group of E89 forms a strong hydrogen bond with the O4’ atom of the ribose-acceptor distance of 2.6 Å and potentially a weaker hydrogen bond with the O2 atom of flavin moiety of FMN with the donor-acceptor distance is 3.3 Å.

**Figure 5.**
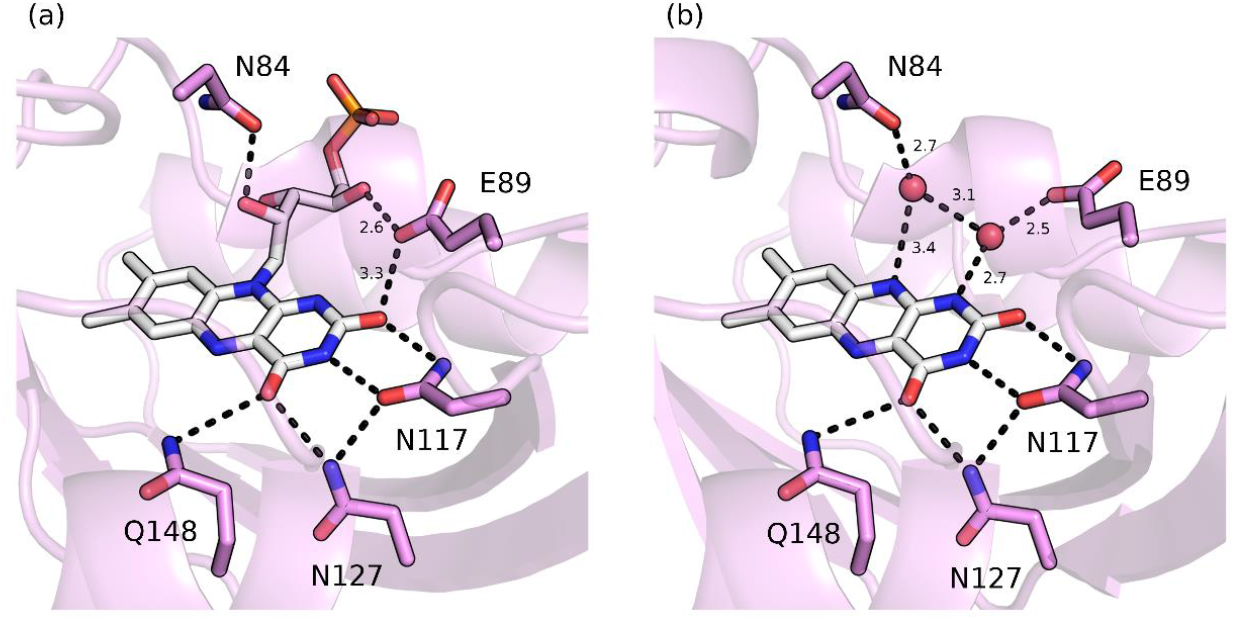
Crystallographic structures of LSSFbFP in complex with FMN**(a)** and LC **(b)**. Ribityl group is replaced with water molecules W1 and W2 in LSSFbFP-LC. Chromophores and key side chains are shown in stick representation. Dashed lines correspond to potential hyodgren bonds, numbers indicate donor to acceptor distances in angstrom (Å).

Structure of LSSFbFP-LC was determined at pH 7.5 and was also almost identical to the structure of the original CagFbFP except for the 89th position and the chromophore. In the place formerly occupied by the phosphorylated ribose tail of FMN, two new water molecules are observed in close proximity of N1 and N10 atoms of LC (thereafter referred to as W1 and W2, Figure 5b). The atomic B factor of oxygen atom of W1 and W2 in A/B protein chains were 23.6/18.2 ^2^Åand 38.7/27.4Å^2^, respectively, indicating that W2 is less ordered compared to W1. Contrary to the LSSFbF-PFMN complex, the side chain of E89 does not interact directly with the flavin moiety, but forms a short 2.5 Å hydrogen bond with W1, and W1 is hydrogen-bonded to the N1 atom of LC. As we show below, the W1 and W2 water molecules play a crucial role in ESPT.

### Water chain dynamics in LSSFbFP

Although the crystal structure revealed the existence of a hydrogen bond chain between lumichrome, W1 and the E89 side chain, precise positions of hydrogen atoms remained unclear. Furthermore, it was not clear whether W1 and W2 would remain in their crystallographic positions in the protein in solution at room temperature. Knowledge of presence and dynamics of hydrogen-bonded chains is vital to understanding the mechanism of ESPT. That is because the hydrogen bond chains are basically the paths by which the hydrogen can escape the N1 atom of lumichrome. To address this, we conducted a 500 ns molecular dynamics (MD) simulation of the LSSFbFP monomer in the complex with LC using the Amber force field (Figure 6).

**Figure 6.**
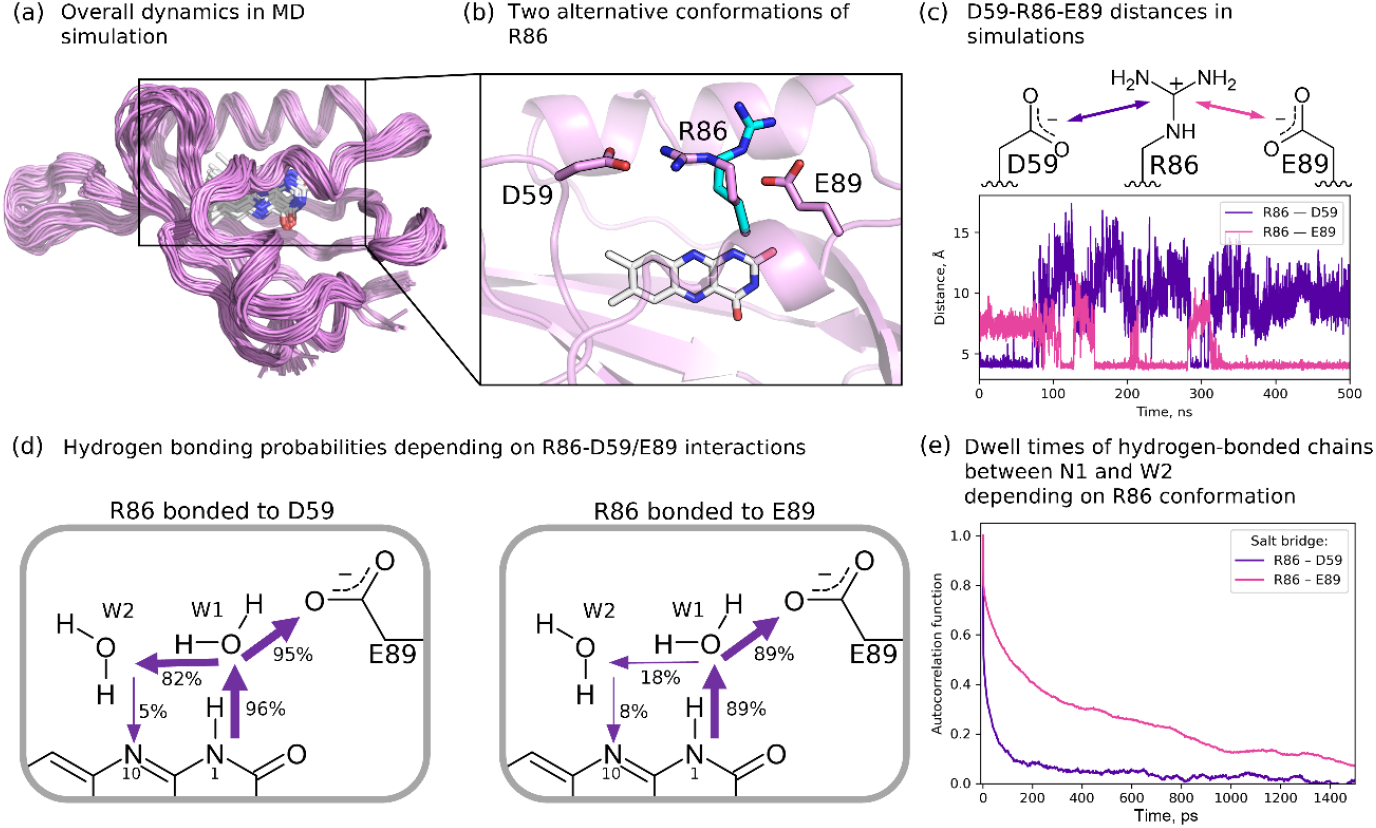
Analysis of molecular dynamics simulations of LSSFbFP in complex with lumichrome**(.a)** Backbone fluctuations in 500 ns MD simulation. **(b)** Two major R86 conformations: bonded to D59 (starting conformation, observed in crystal) and bonded to E8(9**c.)** Distances between R86 C_ζ_ and D59 C_γ_ or E89 C_δ_ observed in MD simulation.**(d)** Presence of hydrogen bonds in MD simulations depending on R86 conformation. R86 was restrained to remain bonded either to D59 or E89**(.e)** Distribution of N1-W2 hydrogen bond dwell times.

In the simulation, we noticed that the R86 residue can reversibly break its salt bridge with the D59 residue and form a salt bridge with the E89 residue instead (Figure 6b,c). The latter conformation of R86 is not observed in the crystal structure, while having a potential to alter the dynamics of water molecules around the lumichrome. We repeated the simulation using the CHARMM36m force field and observed the similar effect (Figure S7). Our theoretical calculation using ASEC FEG showed that this alternative conformation should not affect the absorption spectrum (see Methods and Figure S8).

To simplify the system, we performed two additional 60 ns simulations, where the R86 side chain was softly restrained to remain bonded either to the E89 or to the D59 side chain. To assess the hydrogen bond connectivity of the system, we employed simple geometric criteria (see Methods and Figure S9). Additionally, we estimated the lifetimes of hydrogen-bonded chains using the autocorrelation function. They exhibited a fast decay of a few picoseconds related to rapid fluctuations of hydrogen atoms and a slow decay of > 50 ps related to water molecule reorientation and movement.

In all simulations, W1 remained stably bound to lumichrome (N1→W1 chain). Furthermore, the continuous hydrogen bond chain N1→W1→E89 existed in approximately 90% of simulation time. This indicates that ESPT could potentially happen through deprotonation of LC and protonation of E89 (Mechanism A, Figure 7). Also in all simulations, regardless of the R86 conformation, we observed that the continuous hydrogen bond chain N1→W1→W2→N10 does form for a fraction of time. This indicates that ESPT could happen without deprotonation of LC and result in isoalloxazinic form of LC with protonated N10 (Mechanism B, Figure 7). Finally, a hydrogen atom bound to the N1 atom of LC could just escape to the outer solvent (Mechanism C, Figure 7). This would require at least the N1→W1→W2 chain to be present. Interestingly, the conformation of R86 strongly affects this hydrogen bond chain (Figure 6d). It is mostly present when R86 is bound to D59, and mostly not present when R86 is bound to E89. Also, the lifetime of this chain changes is increased almost tenfold when R86 is bound to E89 (Figure 6e).

**Figure 7.**
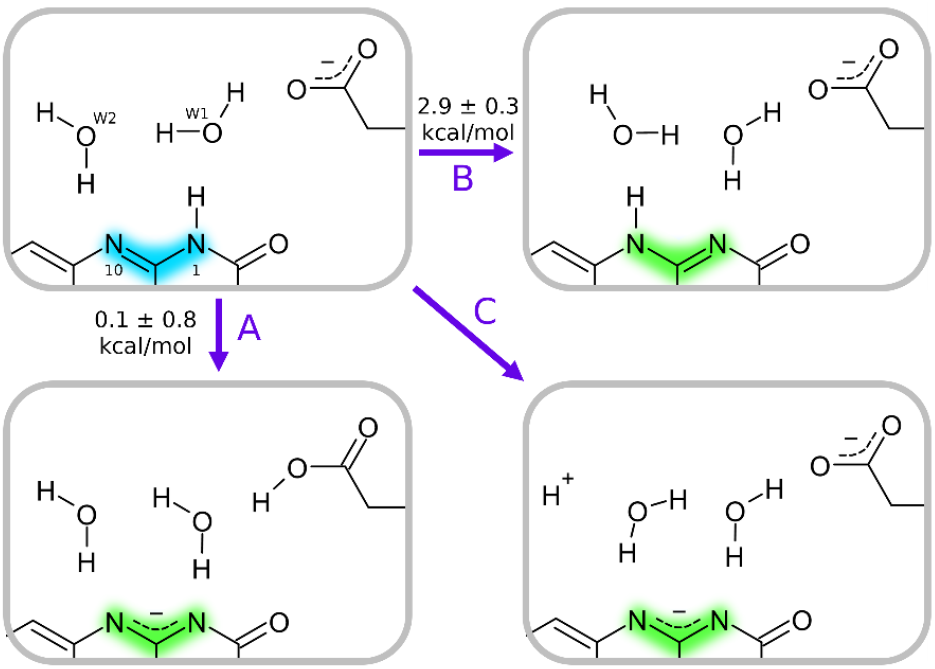
Possible mechanisms of ESPT in LSSFbF-PLC. Colored halos correspond to emission colors of respective compounds.

### Proton transfer mechanism in LSSFbFP-LC

Above, we underlined three mechanisms of ESPT in LSSFbFP that seemed plausible based on the crystallographic structure and the MD simulations. Our next goal was to choose the most probable one. In order to do this, we investigated the mechanisms A and B using three popular density functionals (B3LYP, PBE0, andωB97X). Mechanism C cannot be reliably evaluated using the commonly available computational approaches and resources. We setimated the energy difference between the initial and final states, averaged it between the three methods, and estimated the error as one standard deviation (Figure 7). Based on that, we conclude that while mechanism A is possible, mechanism B is likely energetically disfavored. Also, mechanism B suggests formation of the isoalloxazine form of LC, which, according to our knowledge, has not been convincingly observed for LC.

While mechanism A is possible, it can not explain the nanosecond lifetime and overall complexity of conversion from blue-to green-emitting state as seen in time-resolved spectroscopy (Figure 3). At the same time, mechanism C could explain time-resolved data particularly well. Mechanism C depends on the presence of the N1→W1→W2 chain. While the LC is excited by the photon, the N1→W1→W2 chain might be present or not, and if not, the time of its formation is dependent on the current conformation of the R86 (Figure 8). This would explain the three blue states with the same spectra but different characteristic times of transitions to the green state. Possible state occupancies and corresponding simulated fluorescence properties are presented in Figure S10.

**Figure 8.**
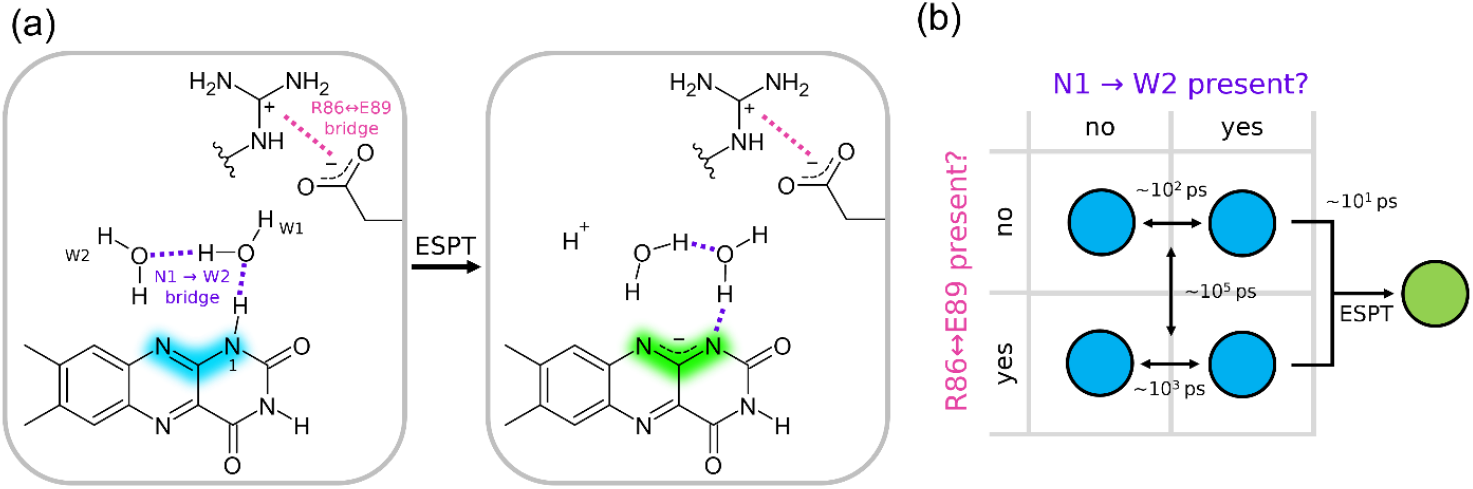
Proposed mechanism of ESPT in LSSFbFP**(.a)** Detailed view of N1→W1→W2 hydrogen bond chain and R86↔ E89 salt bridge before and after ESPT.**(b)** Proposed scheme of post excitation transitions of LC in LSSFbFP. Four blue states are differentiated by presence of hydrogen bond chain and salt bridge.

Finally, we note that the mechanisms A and C could indeed happen simultaneously. Generally, the negatively charged side chain of E89 probably acts as an attraction center that rips the polar hydrogen from the lumichrome. After that, the hydrogen could protonate the side chain or diffuse into the outer solvent, which depends on hydrogen bond connectivity and protein conformation at the moment of excitation.

## Discussion

Flavin-binding proteins remain an exciting area of studies, displaying versatile chemical and light-reactive properties and finding ubiquitous applications as enzymes and potical tools. While studying a model flavoprotein, CagFbFP, we discovered its variant with unique properties: absorption maximum at 390 nm, with photoexcitation resulting in excited state proton transfer, which causes a large apparent Stokes shift. Previously observed photoreactions in flavoproteins included formation of the flavin-(C4a)-cysteinyl covalent bond in wild type LOV domains^26^ and electron transfer to flavin in cryptochromes, photolyases and fatty acid photodecarboxylases^27,28^.

In some scenarios, LSS is a very desirable property of fluorescent tags alongside with brightness, photostability, redshifted emission and pH stability^29 31^. While many popular fluorescent proteins have Stokes shifts below 70 nm, those with LSS, typically over 100 nm, offer several advantages. First, the wider gap between the absorption and emission spectra helps to minimize the inner filter effect, ensuring that fluorescence is not absorbed by the tag itself^32^. Second, LSS enables fluorescence detection without interference from the excitation light, thereby enhancing the signa-lto-background ratio^33^. Finally, proteins with LSS are particularly valuable for multicolor microscopy applications, such as multicolor imaging using a single excitation source^34^ or using a single detection channel and more complex scenarios like observing two distinct FRET pairs within the same sample^35^ or conducting multicolor superresolution microscopy^36^.

Several different mechanisms are responsible for LSS in fluorescent tags used in multicolor microscopy. Many synthetic molecules, which undergo intramolecular charge transfer upon excitation, display LSS due to non-specific solvent effects^37 39^. Additionally, the emission could be shifted from the excitation through specific rapid relaxation of the solvent or amino acid side chains surrounding the excited chromophore. This relaxation process increases the apparent Stokes shift in several genetcially encoded tags, such as mPlum (compared to its predecessor mRFP)^40^, or CagFbFP-I52V-A85Q (compared to the original CagFbFP^2^)^3^. Finally, following excitation, changes in chromophore such as tautomerization or ESPT can lead to the LSS. The latter mechanisms explain the photophysical characteristics of fluorescent proteins exhibiting the largest known Stokes shifts^41^.

To the best of our knowledge, all previously developed genetically encoded LSS fluorescent proteins are based on GFP-like orRFP-like chromophores, whereas LSS tags based on other natural chromophores remain largely unexplored. Overall, photophysics of RFP-like LSS tags closely resembles the photophysics of LSSFbFP-LC studied here. The chromophores found in these proteins contain a phenolic group, whose protonation significantly affects the fluorescence color^42^, similarly to how protonation of N1 effects the color of lumichrome. Additionally, pKa value of both chromophores drops after the excitation^43^. Investigating this mechanism on the exemplary RFP-like tags LSSmKate1 and LSSmKate2^44^ helped in identification of a single mutation that allowed for the rational design of a series of red fluorescent LSS proteins, with emission maxima spanning 560-640 nm and Stokes shifts exceeding 120 nm, all based on RFP variants^45^. This mutation played a similar role to the Q89E mutation discovered in our work. In both cases, the mutation adds a proton acceptor in the form of a carboxylate group into a close proximity to the chromophore. LSSFbFP-LC has a similar excitation and emission spectra to LSS tags based on GFP^35^. To the best of our knowledge, the mechanism behind the large Stokes shifts of GFP-like tags remains unclear due to the absence of an obvious proton acceptor.

Reliance of LSSFbFP on lumichrome as a chromophore is both an advantage and a disadvantage. Existence of different tautomers and redox states with distinct fluorescence colors and wide-ranging optical properties makes lumichrome a promising fluorophore. Photoconversion offers new possibilities for complex imaging regimes, where previously illuminated molecules can be easily distinguished from non-illuminated ones. On the other hand, lumichrome may not be easily available for binding to proteins in all systems *in vivo*. Indeed, newly expressed LSSFbFP primarily forms a complex with common naturally occurring chromophores such as RF, FMN and FAD (although some LC may also be found, Figure S2 and reference^11^). To develop LSS properties, it requires photoconversion to form a complex with LC. Photoconversion may result in undesirable reaction products such as lumiflavin (LF) and its analogs. Therefore, LSSFbFP has room for possible future improvements, including destabilization of its complexes with LF/RF/FMN/FAD and increasing the rate and selectivity of photoconversion to the complex with LC.

Importantly, other FbFPs were previously shown to bind LC: one of them is the original CagFbFP^11^(also studied here), and another is miniSOG, where formation of LC upon photodegradation was unequivocally confirmed using mas-s spectrometry and X-ray crystallography^46^. However, these same time, CagFbFP-LC reveals two very interesting characteristics: (1) notable emission in the indigo/blue range (420-460 nm) at low pH, and (2) pH dependence of absorption and emission spectra with pKa of ~8.5, which potentially could be used for pH detection (Figure 2).

Previously, Kabir and colleagues studied the effects of mutation Q430E of iLOV, which is analogousot mutation Q89E of CagFbFP in our work^47^. Yet, fluorescence spectra of iLOV-Q430E were similar to those of other flavoproteins, and ESPT was not observed. It is not clear at the moment whether flavins bound to iLOV-Q430E could be converted into lumichrome by illumination, and whether the mutation itself accelerates photodegradation.

The last interesting question is whether it is possible to engineer other kinds of LSS tags based on FbFPs with RF/FMN/FAD as fluorophores. To allow for ESPT, the molecule of interest must contain titratable groups, whose protonation or deprotonation should alter the fluorescence emission spectra. While riboflavin is normally neutral, it can also exist in cationic and anionic forms^48^. The cationic form exhibits a blue-shifted absorption, yet unfortunately, neither charged form of riboflavin fluoresces. Alternatively, excitation can result in charge redistribution within the flavin group, leading to specific or nonspecific relaxation of the solvent or protein environment. Consequently, riboflavin demonstrates varying Stokes shifts in different systems^23,49,50^; however, this shift typically does not exceed 90 nm (in free RF in aqueous solution). It remains uncertain whether it is feasible to engineer a protein system with riboflavin displaying Stokes shifts greater than 100 nm. Drawing from the structural similarities, we posit that the same principles apply to FMN and FAD. Consequentlyw, e find it unlikely that FbFPs can be engineered into LSS tags with their natural RF/FMN/FAD chromophore composition.

## Conclusions

Here, we described the discovery of a large Stokes shift flavin-based fluorescent protein, LSSFbFP, which can be obtained *in vivo* and *in vitro*, with absorption maxima at 340-350 and 395-405 nm. We used spectroscopic and structural methods to show that it binds lumichrome as a fluorophore, which undergoes excited state proton transfer upon illumination. Our findings thus present a new type of photoreaction in flavoproteins and expand the range of genetically encoded LSS fluorescent tags beyond the GF-P and RFP-like family.

## Data availability

Crystallographic data for LSSFbFP-LC and LSSFbFP-FMN were deposited into the Protein Data Bank under the accession codes 9U5V and 9U5N, respectively.

## Acknowledgements

We are grateful to SSRF for providing the opportunity to collect crystallographic data at the beamline BL17UM. We are grateful to Dr. Gleb Bourenkov for assistance with the X-ray diffraction data collection at the P14 beamline operated by EMBL Hamburg at the PETRA III storage ring (DESY, Hamburg, Germany). The study was partially supported by the Ministry of Science and Higher Education of the Russian Federation (agreement 0750-3-2025-662, project FSMG-2025-0003).

## Methods

### Random mutagenesis and cloning

Site saturation mutagenesis of position Q89 of the original pET11-CagFbFP plasmid was carried out using a two-step protocol. In the first step, a megaprimer 152 nucleotides long was generated through polymerase chain reaction (PCR) with a forward primer (TATACATATGGCCAGCGGTATGATTGTT) and a Q89X reverse primer (CATCGGTCTGCGGACCMNNCAGAAAACGAGCATTACGAC), which included a randomized MNN codon at the targeted mutagenesis site. The second step involved circular PCR on the original plasmid mixed with the megaprimer. The resulting linear DNA fragments were then transformed into the DH10β *E. coli* strain. Subsequently, the cells were plated on LB-agar Petri dishes supplemented with 150 μg/ml ampicillin and were incubated at approximately 37 °C for 16 hours. All visible colonies were harvested from the agar plates and resuspended in 1 ml of LB medium. Plasmids were extracted from the suspension using an extraction and purification kit (Evrogen, Russia). The resulting plasmids were then transformed into the C41 E. coli strain. The cells were plated on LB-agar Petri dishes supplemented with 1 mM IPTGand 150μg/ml ampicillin, and incubated for 16 hours at 37 °C.

### Clone selection

96 colonies of transformed C41 cells were transferred to a 96-well plate (Greiner, Austria) and resuspended in 100μl of buffer containing 300 mM NaCl, 50 mM Tris-HCl, pH 8.0. Emission spectra of each colony were recorded using Synergy H4 Hybrid Microplate Reader (BioTek, USA). Following the analysis of emission spectra shifts and apparent brightness, four mutants with red-shifted spectra were chosen for protein expression. One of them had the Q89E mutation.

### Expression and purification

Selected cells were cultured in 2 L shaking flasks in 250 ml LB containing 150 mg/ml ampicillin at 37 °C. After reaching OD ~0.6-0.8 expression was induced by adding 1 mM IPTG followed by 5 hours incubation at 37 °C. Additionally, for light illuminated expression, we wrapped the shaking flask with an LED strip emitting cool white light (24 W, color temperature 6500 K). After adding 1 mM IPTG, the flask was incubated for 20 hours at 18 °C.

Harvested cells were resuspended in a buffer containing 300 mM NaCl, 50 mM Tris-HCl, pH 8.0 and disrupted by incubation in a water bath at 95 °C for 10 min followed by 10 min incubation at 0 °C. Cell debris was removed by centrifugation at 10,000 g for 20 min at 10 °C. N-i nitrilotriacetic acid (Ni-NTA) resin (Qiagen, Germany) was added to the supernatant with desired protein and incubated for 1 hour at 4 °C. After that the resin was transferred onto the gravity flow column and washed by buffer containing 300 mM NaCl, 50 mM Tris-HCl, pH 8.0 with 20 mM imidazole to get rid of nonspecific binding. The protein was eluted by a buffer containing 300 mM NaCl, 50 mM Tris-HCl, pH8.0 with 200 mM imidazole.

### Spectroscopy

Steady state excitation spectra were measured using Synergy H4 Hybrid Microplate Reader (BioTek, USA) with emission detected at 520 nm. For steady state emission measurements, samples were excited by a M455F3 LEDat 455 nm (Thorlabs), and the emission spectra were recorded using AvaSpec-2048L spectrometer (Avantes). For absorbance measurements, samples were illuminated by AvaLight-DHc full-range light source (Avantes), and spectra were recorded by the same AvaSpec-2048L spectrometer.

For time resolved fluorescence measurements, a set of fluorescence decay kinetics was measured in a time-correlated single photon mode using a detector with an ultra-low dark count rate (HPM100-07C, Becker&Hickl, Germany) coupled to an ML-44 monochromator (Solar LS, Belarus) used for tuning of the detection wavelength from 400 to 600 nm in 5 nm steps. Fluorescence was excited at 375 nm (repetition rate 80 MHz, pulse width 150 fs, average optical power 1 mW) using the second harmonics ofa femtosecond optical parametric oscillator (TOPOL-1050-C, Avesta Project LTD, Russia) pumped by a femtosecond Yb laser (TEMA-150, Avesta Project LTD, Russia). The emission signal was collected perpendicular to the excitation beam. The temperature of the samples was stabilized at 25C by a thermostatic cuvette holder Qpod 2e with a magnetic stirrer (Quantum Northwest, USA).

For low temperature fluorescence measurement, samples were cooled to 15K using PT403 closed-cycle helium-cooled cryostat (CryoMech, USA). Fluorescence excitation was performed with an ultraviolet diode with an emission range of 360 370 nm. Spectra were recorded using the HR-series spectrometer (Ocean Insight, USA) with a spectral resolution of 1.75 nm. The decay of phosphorescence was recorded using the same experimental setup at 45K. The sample was illuminated with a diode for a few seconds, after which the diode was momentarily switched off while the emission spectra were recorded every 50 ms. The spectra related to the phosphorescence decay were selected based on the absence of fluorescence at 430 nm.

### pH-dependence

For the pH-dependent measurement of emission and absorption spectra, a series of buffers was prepared, each containing 375 mM NaCl and 50 mM of MES, MOPS, HEPES, CAPS, and TRIS. Subsequently, the pH values were adjusted by incrementally adding HCl or NaOH until reaching the final NaCl, 50 mM Tris-HCl at pH 8.0 was mixed in a 1:7 ratio with the prepared buffers. Additionally, we ensured that the addition of the protein solution did not cause deviation of more than 0.1 from the intended pH value.

### Protein characterization

Extinction coefficients were measured as described previously via denaturation in guanidine hydrochlorid^1^e^0^. Quantum yields were measured as described previousl^1^y^0^. Melting curves of proteins were measured using the Rotor-Gene Q realtime PCR cycler (Qiagen, Germany) with excitation at 470 nm or 365 nm and emission measured at 510 nm. Fluorophore dissociation constants were measured exactly as described previously^51^.

### Global analysis of time-resolved spectra

Time resolved fluorescence data was fitted using following functions:

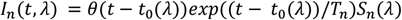

Where *t* is time, *λ* is wavelength, *T*_*n*_ is life-time of temporal component, *S*_*n*_(*λ*) is decay associated spectra, *t*_0_(*λ*) is time of excitation pulse with*λ*-dependent dispersion correction which was fitted as third-degree polynomial,*θ*(*t*) is Heaviside step function and n is the index of the temporal component. The total intensity was assumed to be described by:

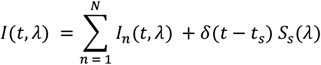

Where last term describes light scattering, *δ* is Dirac delta function and*t*_*s*_, *S*_*s*_(*λ*) are free parameters. Total intensity was further convoluted with instrument response function (IRF) that was measured directly by measuring time resolved Raman scattering of the buffer. More precisely IRF approximated as a sum of two Gaussian functions was used. All parameters were optimized using coordinate descent.

Time resolved phosphorescence data was fitted using following function:

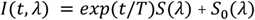

Where *I*(*t, λ*) is total intensity, *S*(*λ*) is decay associated spectrum, *S*_0_(*λ*) is baseline spectrum and T is phosphorescence lifetime. Final spectrum was obtained by averaging calculated *S*(*λ*) from 42 independent experiments.

### Crystallization, data collection, and structure determination

Concentrated LSSFbFP-LC was crystallized using a sitting drop vapor diffusion approach using the NT8 robotic system (Formulatrix, USA). The drops contained 150 nL concentrated protein and 150 nL reservoir solution. Crystallization plates were stored at 20 °C. Elongated crystals reaching 100 400 µm appeared after 1 week in the wells with the precipitant solution containing 0.2 M Sodium formate, 0.1 M Bi-sTris propane Buffer pH 7.5 and 20 % w/v PEG 3350. The crystals were harvested using micromounts, cryoprotected by immersing into 20 % glycerol, flash-cooled and stored in liquid nitrogen. Diffraction data were collected at the BL17 beamline of the Shanghai Synchrotron Radiation Facility (SSRF). Data integration was performed using XD^5^S^2^, and scaling was completed using the STARANISO web serve^53^r, revealing anisotropic diffraction limits of 1.635, 1.760 and 2.042 (Table S1). Elliptically truncated data were subsequently utilized for molecular replacement in MOLREP^54^, using the PDB ID 6RHF mode^2^l^2^ as the template. Structure refinement was conducted in Refmac5^55^, with related statistics provided in Table S2.

Concentrated LSSFbFP-FMN was crystallized using a sitting drop vapor diffusion approach using the Mosquito robotic system (SPT Labtech, United Kingdom). The drops contained 100 nL concentrated protein and 200 nL reservoir solution. Crystallization plates were stored at 20 °C. Elongated crystals reaching 200 µm appeared after 1 week in the wells with the precipitant solution containing 20% v/v ethylene glycol, 10% w/v PEG 80000, .02 M sodium formate, 0.02 M ammonium acetate, 0.02 M sodium citrate tribasic dihydrate, 0.02 M potassium sodium tartrate tetrahydrate, 0.02 M sodium oxamate, 0.1 M imidazole and 0.1 M MES monohydrate pH 6.5. The crystals were harvested using micromounts, cryoprotected by immersing into 20 % glycerol, flash-cooled and stored in liquid nitrogen. Diffraction data were collected at the P14 beamline of the PETRA III at Deutsches Elektronen-Synchrotron (DESY). Data integration was performed using XDS, and scaling was completed using AIMLESS^56^. The data were subsequently utilized for molecular replacement in MOLREP, using the PDB ID 6RHF model as the template. Structure refinement was conducted in Refmac5, with related statistics provided in Table S2.

### Molecular dynamics simulation

Molecular dynamics simulations were conducted using OpenMM 7.6^57^. Initial coordinates for modeling were taken from chain B of crystallographic structure of LSSFbFP in complex with lumichrome. Protonated structure of monomeric protein was prepared using CHARMM-GUI PDB Reader & Manipulator too^5^l^8 60^. Protonation states of titratable residues were assigned to pH = 8 based on pKa. The lumichrome was modeled in its neutral aloxazinic form. The system was neutralized by adding Na+ and C-l ions to ionic strength of 0.15M. The protein was solvated in a cubic OP^6^C^1^ water box with a water layer of 10 Å around the protein using CHARMM-GUI Solution Builde^6^r^2,63^. Crystal waters were kept. Amber ff19SB^64^ parameters were used for protein and GAFF2^65,66^ for lumichrome. Additionally, we calculated accurate RESP charges for lumichrome using the following procedure. Ground state geometry of neutral aloxazinic lumichrome was optimized using B3LYP^67 70^ density functional with ma-def2-SVP^71,72^basis set and D3^73^ dispersion correction. After that, electron density was calculated using B3LYP density functional and 6-311G*^74^* basis set. DFT calculations were performed using ORCA^75 77^ and RESP chargers were calculated using Multiwfn^78,79^. RESP charges were used instead of ones provided by CHARMM-GUI. System was energy minimized using the steepest descent method. Then, 1 ns of equilibration were performed. During the equilibration and minimization steps harmonic restraints were applied to backbone atoms of the protein and carbon atoms of the ligand. Force constant of restraints was 5 kcal/mol Å^2^ during the minimization and was gradually reduced to zero during equilibration. Finally, a production run was carried out for the desired amount of simulation time. Langevin middle integrator^80^ with a time step of 2 fs for short runs (less than 100 ns) and 4 fs for long run (500 ns), friction coefficient of 1 ps and reference temperature of 300K was used. Pressure was kept by Monte-Carlo barosta^8^t^1,82^ at 1 bar, which was applied every 100 steps. Cutoff of 1 nm were implemented for calculating Lennard-Jones interaction and particle mesh Ewald method^83^ was used for long-range electrostatic interactions with error tolerance of 0.0005. Length of bonds containing hydrogen were constrained during the simulation. Hydrogen mass repatriation was used for 500 ns run, hydrogen mass was set to 1.5 Da. For simulation with restrained behavior of R86 position we employed harmonic flat bottom potential between carbon atom of carboxylate group of D59 or E89 and carbon atom of guanidinium group of R86. The potential had the following form:

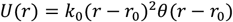

Where r is a distance,*θ* is Heaviside step function and *k*_0_ was set to 2 kcal/mol Å^2^ and *r*_0_ to 4.8 A. Constrained simulation where the salt bridge is present between R86 and E89 was started from a frame from an unconstrained simulation in which the bridge was present.

For hydrogen bonds analysis we firstly identified the closest waters to lumichrome at each frame of the simulation. Water 1 (W1) was defined as the water whose oxygen atom is closest to the hydrogen atom bonded to whose hydrogen atom is closest to the N10 atom of lumichrome and not W1. As hydrogen bond criteria we chose the following one. Let D, A and H be donor (heavy atom), acceptor (heavy atom) and donated hydrogen atom. Two molecules were considered hydrogen bonded if D-A distance is less than 3.4 A and angle H-D-A is less than 35 degree. Numbers were chosen artificially and strictly speaking, the criterion speaks more about the geometric possibility of hydrogen bond formation rather than its actual presence. Analysis was conducted using MDTraj python library^84^.

### QM/MM modeling

Initial coordinates for protein heavy atoms and lumichrome were taken from chain B of crystallographic structure of LSSFbFP. Solvent box and hydrogens were added to the system using CHARMM-GUI Solution Builder. Added solvent was equilibrated around the protein and crystal water molecules using OpenMM 8.0.1During the equilibration initial atoms including crystallographic water were hardly restrained to their crystallographic positions. Equilibrated system was energy minimized for further modeling. System states with occurred proton transfer were prepared by manually moving hydrogen atoms in PyMOL.

QM/MM modeling was conducted using ORCA 5.0.3 Additive QM/MM scheme with electrostatic embedding was applied. Prepared structure was divided into two subsystems: QM subsystem containing lumichrome, E89 sidechain and 5 relevant water molecules. For the QM part of the QM/MM calculation, B3LYP density functional for ground state and time-dependent B3LYP^69,85^ for excited state with def2-SVP^71^ basis set and D3dispersion correction were used. Bonded interaction between subsystems was managed using a link hydrogen atom and charge-shift scheme. QM/MM-optimized geometry were obtained for both ground and first singlet excited state of lumichrome. All MM atoms were fixed during the optimization. After that, ground and excited state energies of obtained geometries was calculated using B3LYP and PBE0 density functional^8^s^6^ with D3 dispersion correction and ωB97X-D3^87,88^ density functional using def2-TZVP^71^basis set.

The ASEC-FEG^89^ method was used to predict the effect of R86 conformation on the spectrum shift. MD steps were conducted using OpenMM and QM steps using ORCA. The MD system was prepared as described above, with additional flat-bottom restraints to maintain desired side-chain orientations. The MD steps consisted of 0.5 ns equilibration and 50 ns production run, and 20 frames were used for ASEC construction. The QM part consisted of one lumichrome molecule. For the QM steps, the B3LYP functional with D3 correction and def2-SVP basis set was used was used for geometry optimization, and the (TD-) B3LYP density functional with the def2-TZVP basis set was used for calculating vertical excitation energies. CHELPG was used for partial charges calculation. The vertical excitation energy converged after only one iteration in all calculations due to the rigidity of the flavin moiety. The final value was obtained by averaging vertical excitation energies from six subsequent iterations.(Figure S8).

