## Supplementary materials for "Discovery of excited state proton transfer in flavin-based fluorescent protein with large Stokes shift"

**Supplementary Data**

**Supplementary Tables 1-2**

**Supplementary Figures 1-11**

**Supplementary Data**

Nucleotide sequences of the studied proteins:

>CagFbFP
ATGGCCAGCGGTATGATTGTTACCGATGCCGGTGCAGATCAGCCGATTGTTTTTGTTAATCG
TGCATTTAGCACCATCACCGGCTATGCACCGAATGAAGTTCTGGGTCGTAATGCTCGTTTTC
TGCAAGGTCCGCAGACCGATGCAGCAACCGTTGCACGTCTGCGTGAAGCCATTGCAGCAGCA
CGTCCGATTCAAGAACGTATTCTGAATTATCGTAAAGATGGTCAGCCGTTTTGGAATCAGCT
GAGCATTAGTCCGGTTCGTGATGAAACCGGCAATGTTGTTGCATTTGTTGGTGTTCAGACAG
ATGTTACCGCACATCATCATCACCATCACTAA

>LSSFbFP
ATGGCCAGCGGTATGATTGTTACCGATGCCGGTGCAGATCAGCCGATTGTTTTTGTTAATCG
TGCATTTAGCACCATCACCGGCTATGCACCGAATGAAGTTCTGGGTCGTAATGCTCGTTTTC
TGGAGGGTCCGCAGACCGATGCAGCAACCGTTGCACGTCTGCGTGAAGCCATTGCAGCAGCA
CGTCCGATTCAAGAACGTATTCTGAATTATCGTAAAGATGGTCAGCCGTTTTGGAATCAGCT
GAGCATTAGTCCGGTTCGTGATGAAACCGGCAATGTTGTTGCATTTGTTGGTGTTCAGACAG
ATGTTACCGCACATCATCATCACCATCACTAA

Amino acid sequences of the studied proteins:

>CagFbFP
MASGMIVTDAGADQPIVFVNRAFSTITGYAPNEVLGRNARFLQGPQTDAATVARLREAIAAA
RPIQERILNYRKDGQPFWNQLSISPVRDETGNVVAFVGVQTDVTAHHHHHH

>LSSFbFP
MASGMIVTDAGADQPIVFVNRAFSTITGYAPNEVLGRNARFLEGPQTDAATVARLREAIAAA
RPIQERILNYRKDGQPFWNQLSISPVRDETGNVVAFVGVQTDVTAHHHHHH

**Table S1: Information about the diffraction anisotropy for LSSFbFP-LC crystals.**

Diffraction limits (Å) and corresponding principal axes of the ellipsoid fitted to the diffraction cut-off surface as direction cosines in the orthogonal basis (standard PDB convention), and in terms of reciprocal unit-cell vectors.

| | Diffraction limits (Å) | Principal axes in the orthogonal basis | Principal axes reciprocal unit-cell vectors |
|---|---|---|---|
| Diffraction limit #1 | 1.635 | ( 1.0000, 0.0000, 0.0000) | a* |
| Diffraction limit #2 | 1.760 | ( 0.0000, 1.0000, 0.0000) | b* |
| Diffraction limit #3 | 2.042 | ( 0.0000, 0.0000, 1.0000) | c* |

Eigenvalues of the overall anisotropy tensor on |F|s ($Å^2$), the same eigenvalues after subtraction of the smallest eigenvalue (as used in the anisotropy correction), and corresponding eigenvectors of the overall anisotropy tensor as direction cosines in the orthogonal basis (standard PDB convention), and in terms of reciprocal unit-cell vectors:

| | Eigenvalues ($Å^2$) | Eigenvalues used in the anisotropy correction ($Å^2$) | Eigenvectors in the orthogonal basis | Eigenvectors in reciprocal unit-cell vectors |
|---|---|---|---|---|
| Eigenvalue #1 | 16.01 | 0.41 | ( 1.0000, 0.0000, 0.0000) | a* |
| Eigenvalue #2 | 15.60 | 0.00 | ( 0.0000, 1.0000, 0.0000) | b* |
| Eigenvalue #3 | 24.19 | 8.59 | ( 0.0000, 0.0000, 1.0000) | c* |

**Table S2. Crystallographic data collection and refinement statistics.**

| Protein | LSSFbFP-LC | LSSFbFP-FMN |
|---|---|---|
| Synchrotron and beamline | SSRF, BL17UM | PETRA III, P14 |
| PDB ID | 9U5V | 9U5N |
| Wavelength, Å | 0.97919 | 0.6888 |
| Resolution range*, Å | 48.099 - 1.663 (1.825 - 1.663) | 48.24 - 1.27 (1.303 - 1.270) |
| Space group | P $2_1$ $2_1$ 2 | P $2_1$ $2_1$ 2 |
| Unit cell | 53.75 Å 107.58 Å 38.93 Å<br>90° 90° 90° | 53.72 Å 109.60 Å 39.05 Å<br>90° 90° 90° |
| Total reflections | 143,157 (6,435) | 1,661,470 (81,927) |
| Unique reflections | 19,963 (999) | 61,767 (2,990) |
| Multiplicity | 7.2 (6.4) | 26.9 (27.4) |
| Completeness (spherical), % | 72.8 (15.3) | 100.0 (100.0) |
| Completeness (ellipsoidal), % | 92.7 (53.1) | - |
| Mean I/σ(I) | 7.9 (1.4) | 14.1(0.6) |
| Wilson B-factor*, Å$^2$ | 19.4 | 19.5 |
| R-merge | 0.173 (1.253) | 0.115 (6.213) |
| R-meas | 0.186 (1.361) | 0.118 (6.329) |
| R-pim | 0.068 (0.521) | 0.023 (1.198) |
| CC1/2 | 0.996 (0.675) | 1.000 (0.432) |
| Resolution range used in the refinement, Å | 48.099 - 1.635, 1.760, 2.042 | 48.24 - 1.27 |
| Reflections used in refinement | 17,359 | 61,752 |
| Reflections used for R-free | 856 | 3,051 |
| R-work | 0.189 | 0.1603 |
| R-free | 0.238 | 0.1893 |
| Number of non-hydrogen atoms | 1,861 | 1,964 |
| macromolecules | 1,657 | 1,676 |
| ligands | 36 | 74 |
| solvent | 168 | 214 |
| Protein residues | 211 | 211 |
| $RMS_{bonds}$, Å | 0.002 | 0.004 |
| $RMS_{angles}$, ° | 0.713 | 1.169 |
| Ramachandran favored, % | 98 | 99 |
| Ramachandran allowed, % | 2 | 1 |
| Ramachandran outliers, % | 0 | 0 |
| Rotamer outliers, % | 0 | 0 |
| Clashscore | 6 | 2 |
| Average B-factor, Å$^2$ | 20.1 | 21.7 |
| macromolecules | 19.8 | 20.5 |
| ligands | 15.4 | 19.2 |
| solvent | 24.6 | 32.0 |

Statistics for the highest-resolution shell are shown in parentheses.
* - anisotropy information is presented in Table S1.

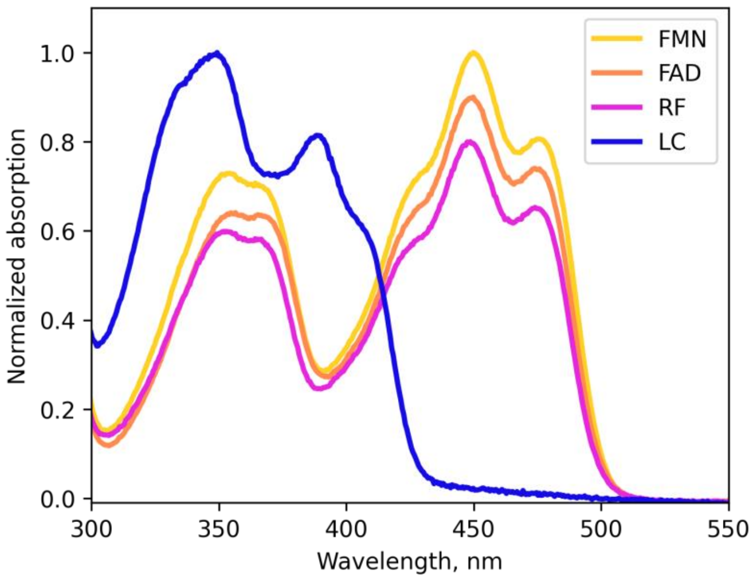


**Figure S1.** Absorption spectra of LSSFbFP reconstituted with different chromophores measured at pH 8. Spectra are normalized to have the absorbance maximum of 1 (FMN and LC), 0.9 (FAD), and 0.8 (RF) for clarity of the presentation.

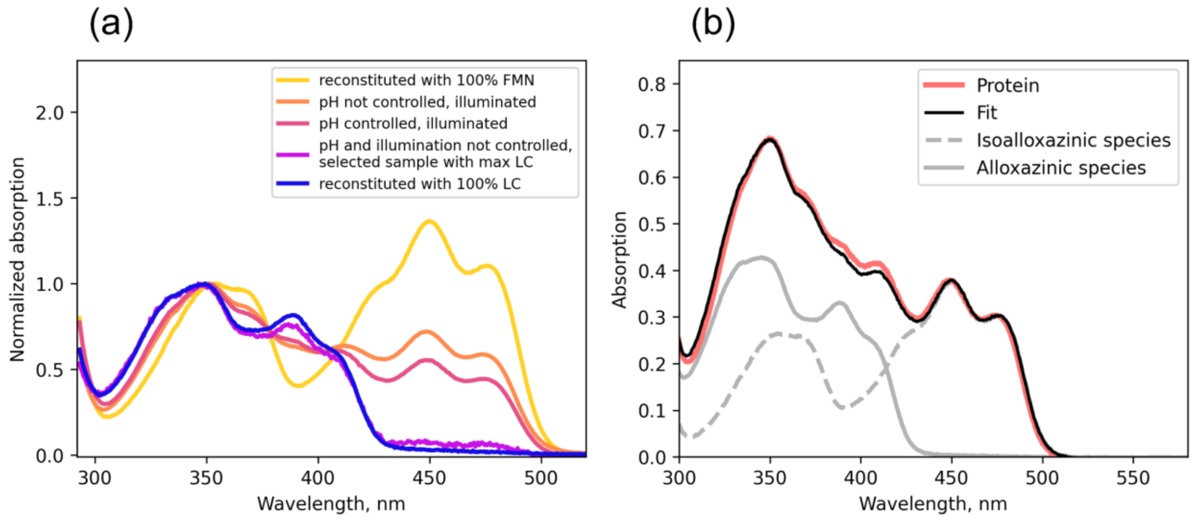


**Figure S2.** LSSFbFP chromophore composition depends on cell culture conditions during expression. (a) Dependence of absorption spectrum of LSSFbFP on cell culture cultivation conditions. (b) Absorption spectrum of LSSFbFP expressed in *E. coli* with pH and illumination controlled. The spectrum is fitted with a weighted sum of spectra of LSSFbFP-LC and LSSFbFP-FMN in order to calculate chromophore composition: 65% LC and 35% FMN.

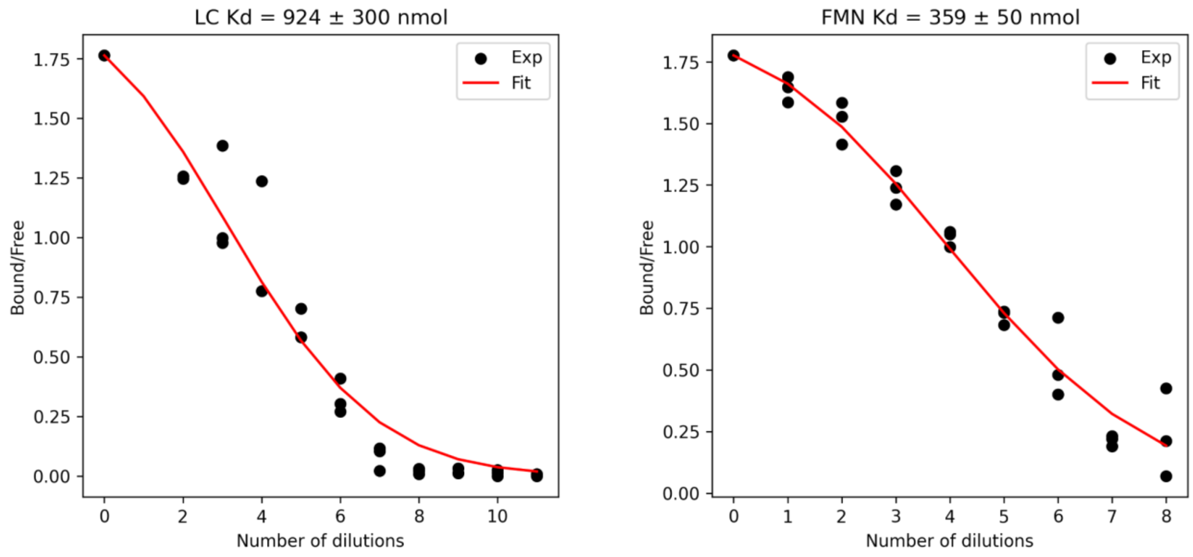


**Figure S3.** Determination of chromophore dissociation constants for LSSFbFP by dilution. Ratio of bound chromophore to free chromophore is determined by linear decomposition of fluorescence spectra. Data from three replicate experiments were fitted simultaneously.

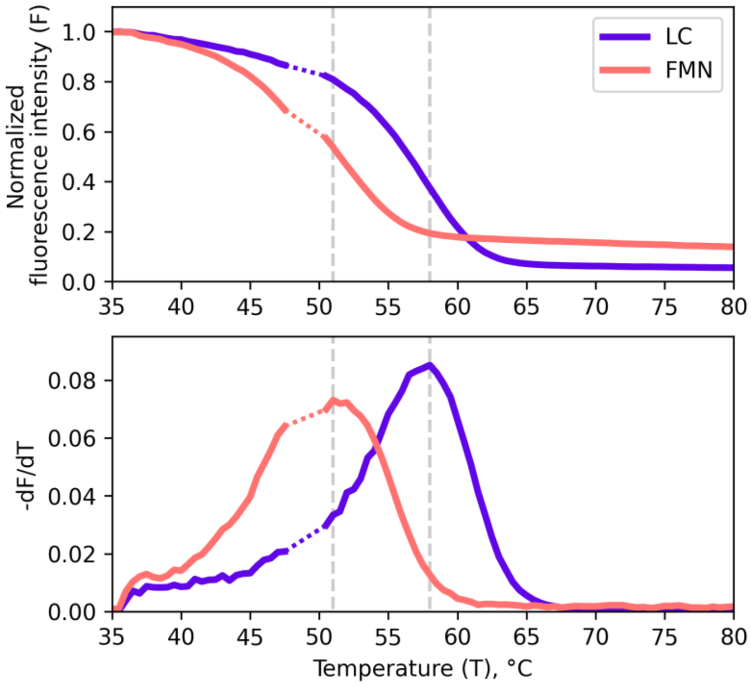


**Figure S4.** Decay of fluorescence of LSSFbFP complexed with lumichrome (LC) and flavin mononucleotide (FMN) upon thermal denaturation. Fluorescence was excited at 365 and 470 nm for complexes with LC and FMN respectively and fluorescence intensity was registered at 510 nm. Data between 48 °C and 50 °C was not measured due to instrument limitations.

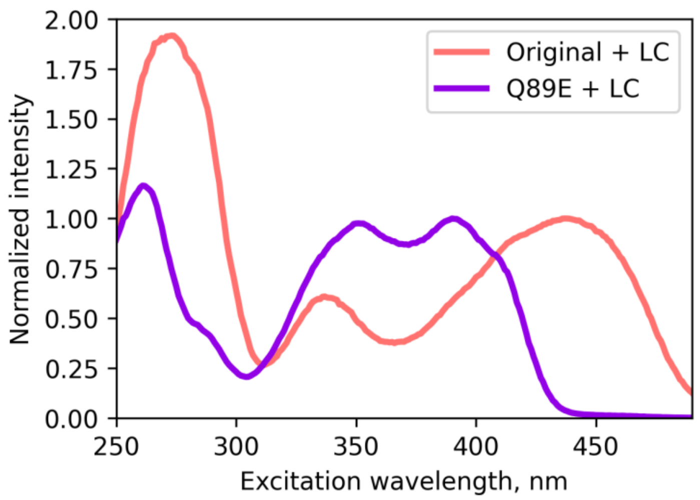


**Figure S5.** Fluorescence excitation spectra of CagFbFP-LC and LSSFbFP-LC at pH 8. Emission is detected at 520 nm. Absorption and emission spectra may be found in Figure 2.

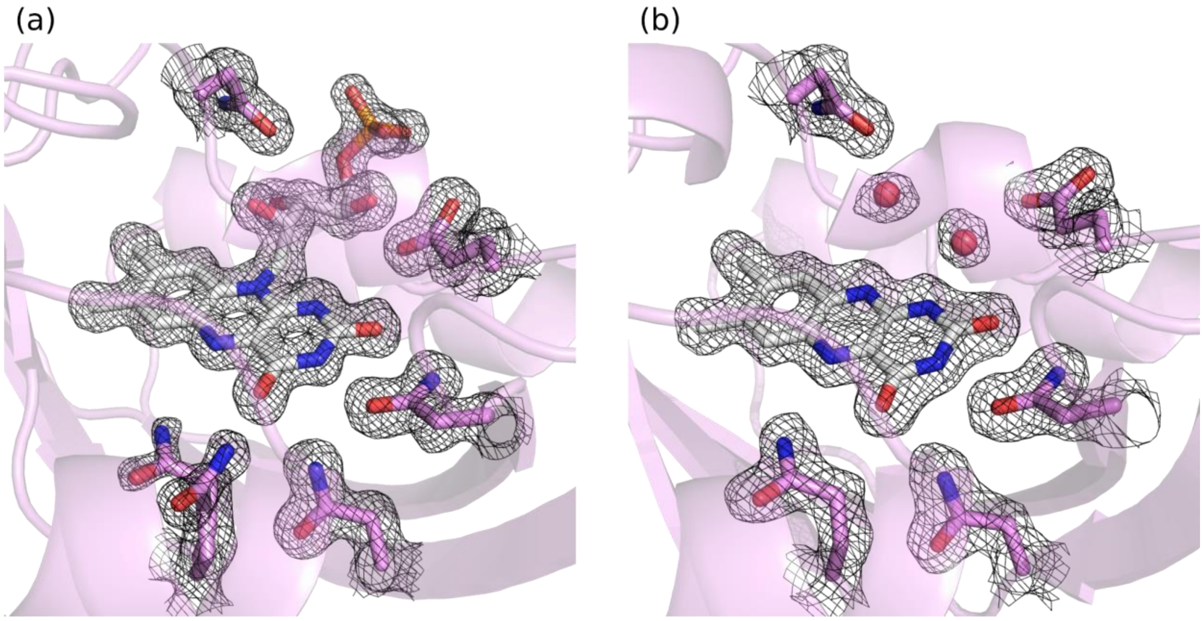


**Figure S6.** Electron densities in the chromophore region for LSSFbFP-FMN (a) and LSSFbFP-LC (b). $2F_o - F_c$ maps are shown at the level of 1 σ.

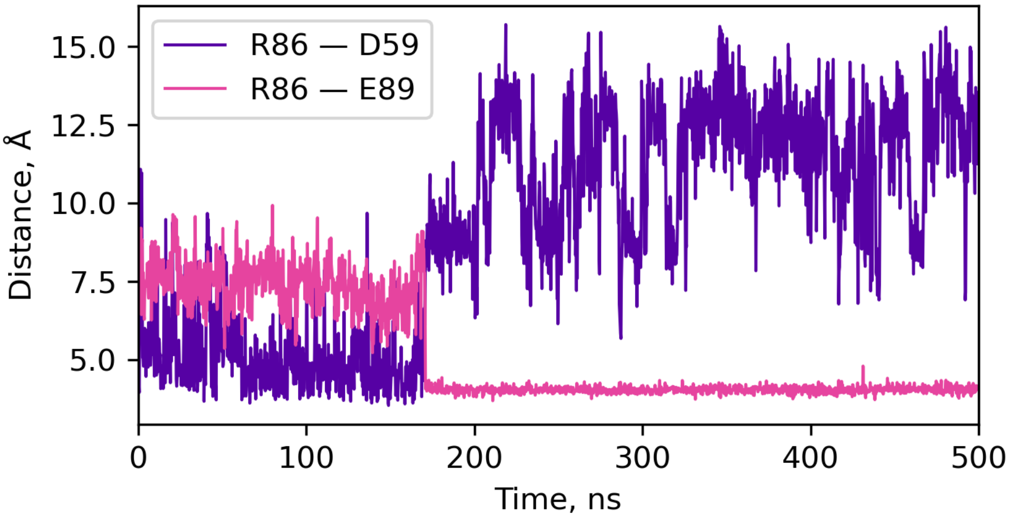


**Figure S7.** Breakage of salt bridge between Arg86 and Asp59 and formation of salt bridge between Arg86 and Glu89 as seen in molecular dynamics simulation employing CHARMM36m force field. Distances are defined as described in the main text.

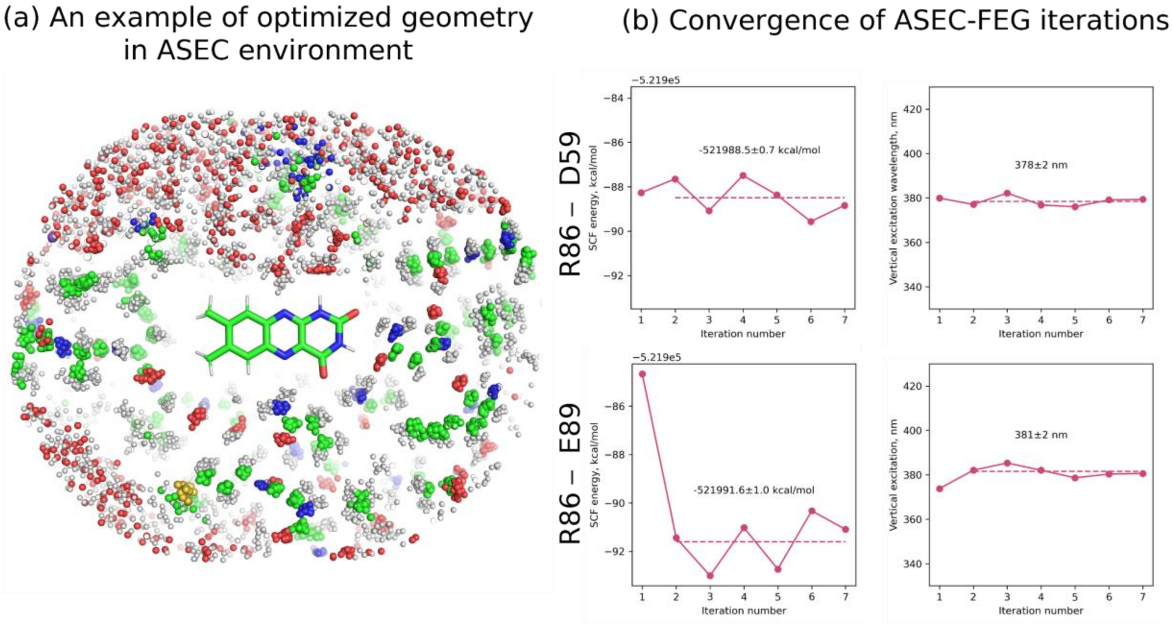


**Figure S8.** ASEC–FEG modeling of lumichrome properties. (a) Optimized geometry in the ASEC environment. (b) Vertical excitation of LSSFbFP-LC calculated using ASEC-FEG iterations. Spectrum shift caused by R86 flipping is negligible.

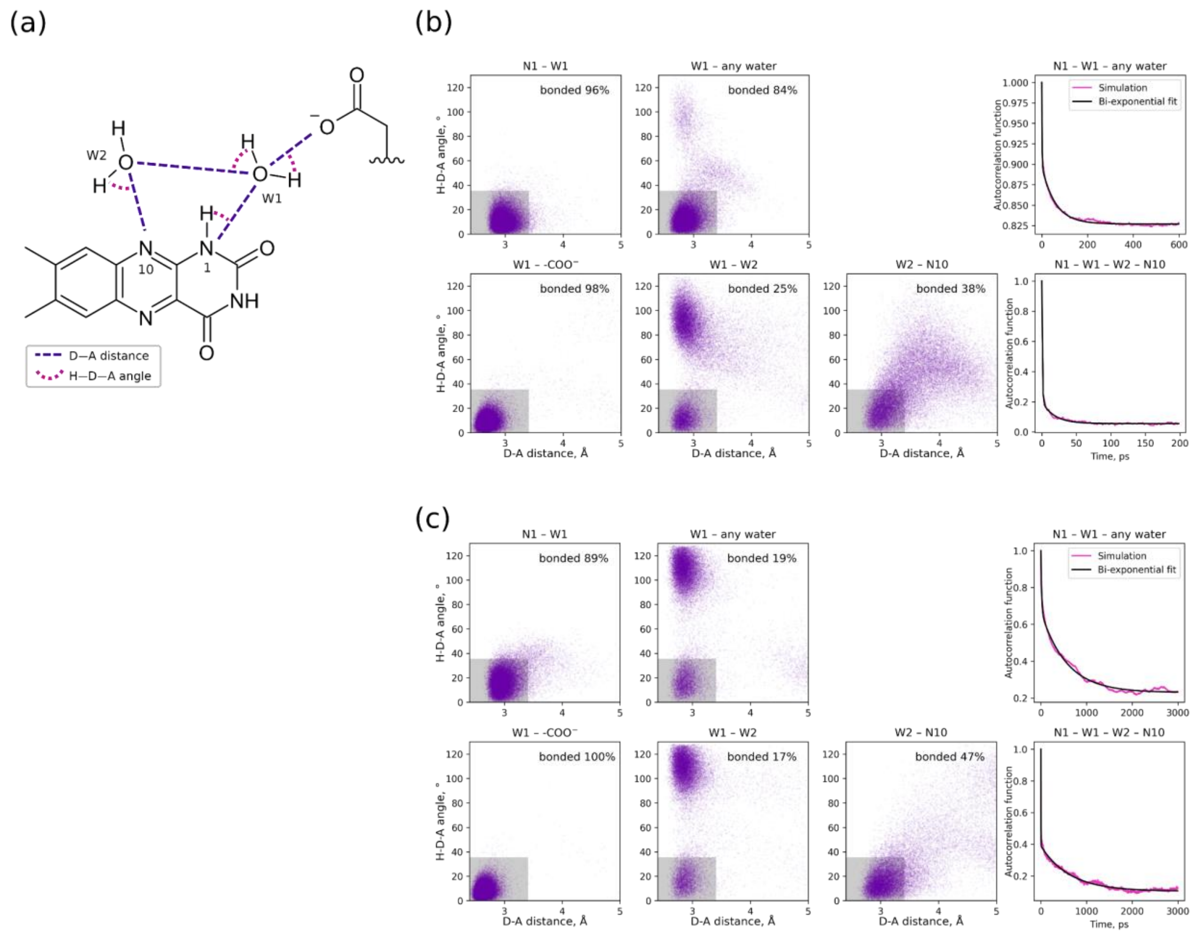


**Figure S9.** Analysis of hydrogen bond patterns in LSSFbFP-LC in MD simulations depending on salt bridges between R86 and D59 or E89. (a) Bond lengths (donor-acceptor distances) and characteristic angles needed to define geometric criteria of hydrogen bond formation. (b) Data for the simulation with R86 restrained to remain bonded to D59. (c) Data for the simulation with R86 restrained to remain bonded to E89. Bonds and angles for several possible hydrogen bonds are analyzed. Distributions of dwell times for hydrogen bond chains N1 - W1 - any water and N1 - W1 - W2 - N10.

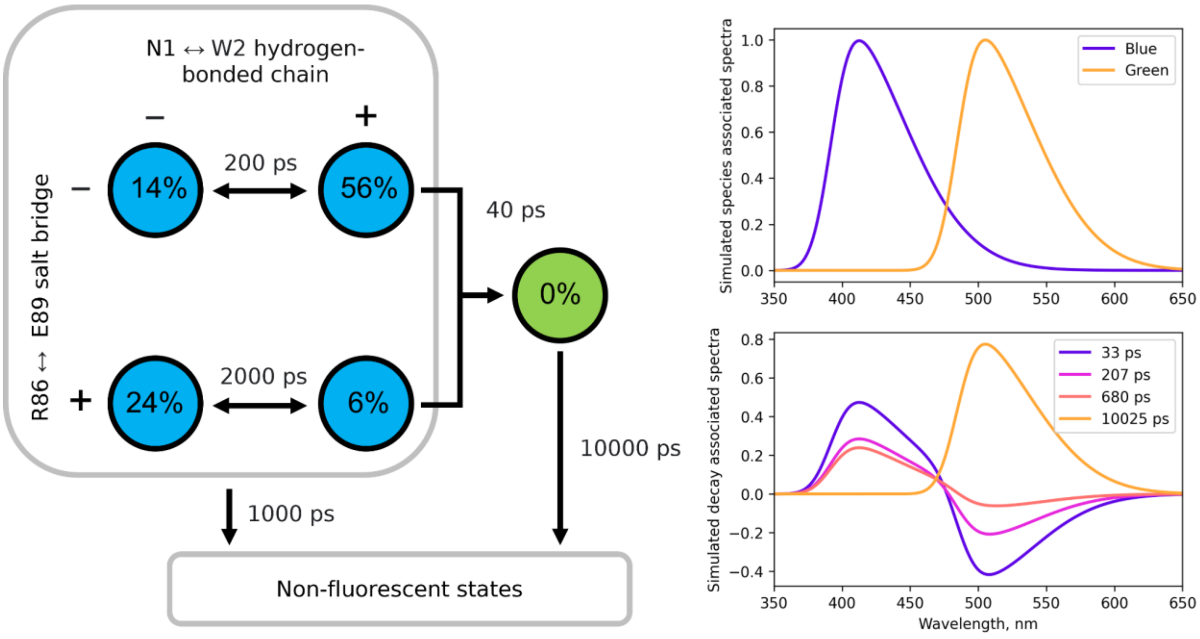


**Figure S10.** Possible scheme of post-excitation transitions of LC in LSSFbFP and corresponding simulated time-resolved fluorescence data. Experimental data is provided in Figure 3g, h. Percentages in the circles represent the populations of the states immediately after excitation, chosen to correspond roughly to the populations seen in the molecular dynamics simulations. Green-emitting state is not populated immediately after excitation.

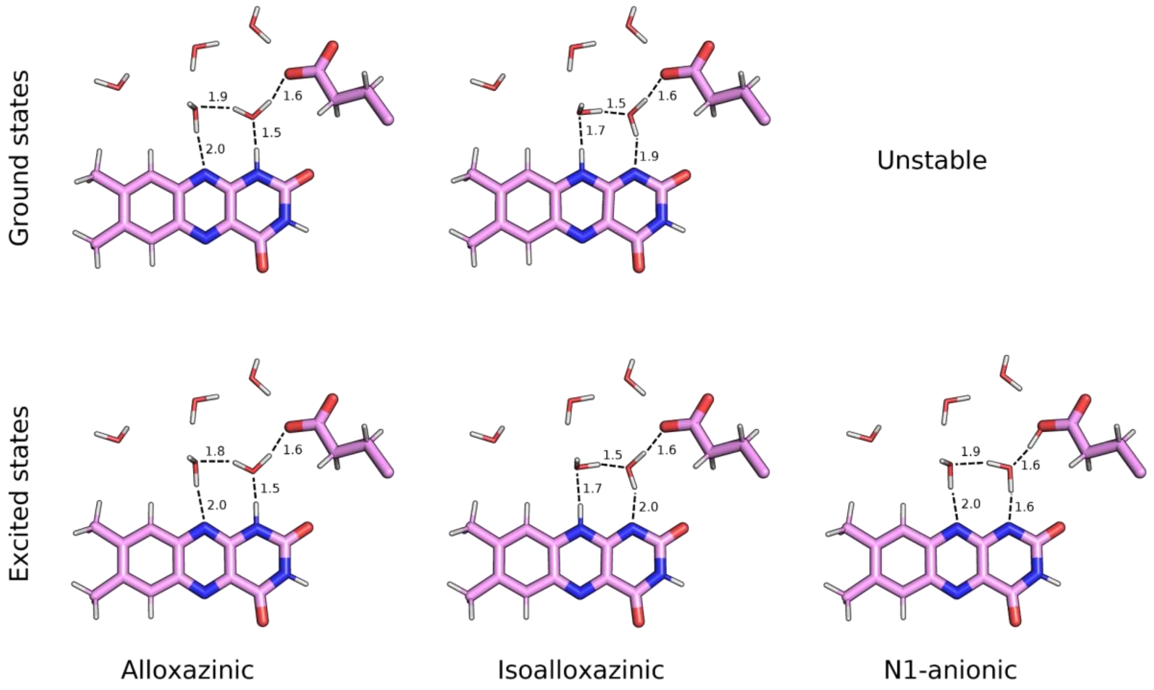


**Figure S11.** Optimized geometries for ground and excited of differently protonated lumichrome. The ground state of N1-anionic lumichrome is unstable.
